# 150 years of research on Vierordt’s law - Fechner’s fault?

**DOI:** 10.1101/450726

**Authors:** Stefan Glasauer, Zhuanghua Shi

## Abstract

150 years ago, the physiologist Karl Vierordt discovered one type of systematic errors in time perception – an overestimation of long durations and underestimation of short durations, now known as Vierordt’s law. Here we review the original study in its historical context and ask whether Vierordt’s law is a result of an unnatural experimental randomization protocol. Using iterative Bayesian updating, we simulated the original results with astonishing accuracy. To validate whether the experimental protocol matters, we compared duration reproduction from two sequences with the same sampled distribution and found that trial-wise variation determines the reproduction error. We conclude that Vierordt’s law is caused by an unnatural yet widely used experimental protocol.

**One Sentence Summary:** Probabilistic modeling reveals that a classical perceptual error - Vierordt’s law - is caused by an unnatural yet widely used experimental protocol.

---

Exactly 150 years ago, in 1868, Karl Vierordt, professor of physiology at the University of Tübingen, published his book “Der Zeitsinn nach Versuchen” (The sense of time according to experiments, *1*), just a few years after Gustav Theodor Fechner’s groundbreaking book “Elemente der Psychophysik” (*2*) and one year after Hermann Helmholtz’s “Handbuch der physiologischen Optik” (*3*). His seminal book was the first quantitative attempt to investigate time perception with the methodology proposed and invented by researchers such as Ernst Weber, Gustav Theodor Fechner, and others.

One of his main findings, and the one that best survived time, is now known as Vierordt’s law. According to this law, short temporal durations tend to be overestimated, whereas long durations tend to be underestimated. Somewhere in between there is an “indifference point” where perceived time is veridical. The mechanisms underlying Vierordt’s law have long remained obscure. Vierordt’s law was considered as an unexplained problem that “currently defies any coherent theoretical treatment” up until 10 years ago (*4*).

For his main experiments, most of them done by Karl Vierordt as an only participant with the help of his assistant. His assistant produced a time interval by two clicks, and he replicated this stimulus by clicking a third time so that the interval between the second and third clicks was perceived as long as the interval between the first and second clicks. The machinery used for the experiments was sophisticated enough to allow recording of stimulus and response from durations below 250 ms to several seconds. Luckily, Vierordt explained his methods in detail and also published most of his data as tables. In the following, we concentrate on his Table A as an example (*1*, p36). It lists the average stimulus duration together with the signed error of reproduction for 22 intervals (from below 250 ms to above 8 s) and the corresponding number of repetitions (ranging from 25 to 83). The whole experiment consisted of overall 1104 trials presented consecutively, which clearly demonstrated the main feature of Vierordt’s law, an overestimation of the short intervals and underestimation of the long intervals with the indifference point around 2.25 s (see Figure 1A).

**Fig. 1.**
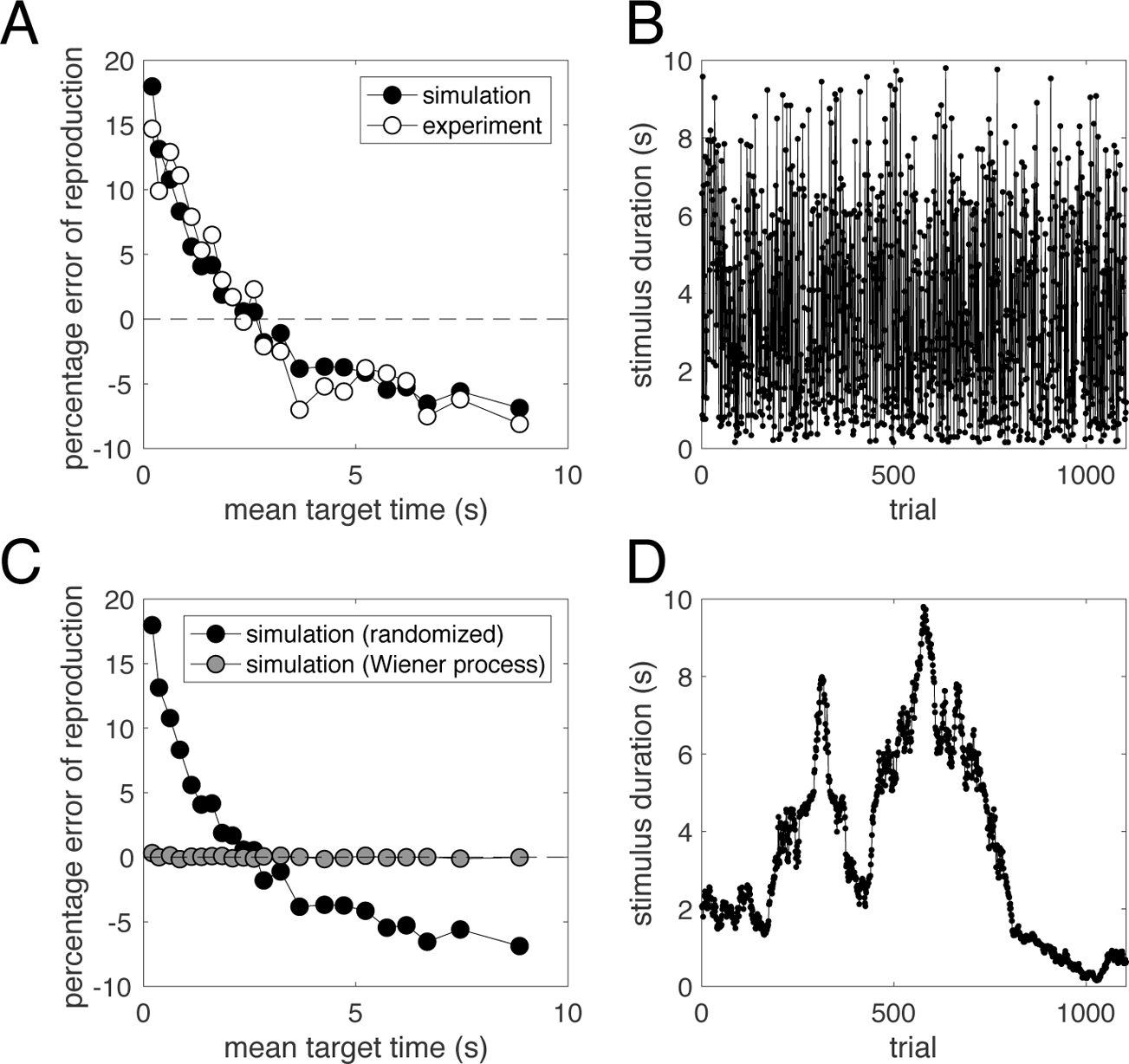
Reproduction data of Vierordt’s durations and iterative Bayesian models. **(A)** data from Vierordt’s original experiment (open circles) and the best fitting model simulation (filled circles). **(B)** The sequence of the durations used for the simulation in **A**. **(C)** Comparison of simulation in **A** (filled circles) with the simulation from the sequence conforming to a random walk or Wiener process (gray filled circles). **(D)** The sequence of the durations for the simulation in **C**. Note that sampled durations in B and D are exactly the same, except for the sequential order.

The method used by Vierordt (*1*, p22) was the “method of average error” that Fechner invented (*2*, Vol. 1, p120 ff, Vol. 2, p148 ff and p343 ff), which Müller (*5*) later referred to as the “method of reproduction” in 1904, now also known as method of adjustment. The method works as follows: a stimulus magnitude N (“Normalreiz”, normal stimulus) is presented and followed by a test stimulus F (“Fehlreiz”, error stimulus), which is adjusted by the participant, so that N and F are perceived as equal. Then the stimulus N is given again, followed by the adjustable F, and so on (*2*, p190 ff). In Vol. 2 (*2*, p343 ff) Fechner explained his method of average error in more detail. He applied 10 measurements of exactly the same condition (and same magnitude) consecutively. If there were multiple magnitudes, the magnitudes were tested in either increasing or decreasing order, and each magnitude was test in a chunk of 10 measurements. Fechner was very accurate about his method: for example, concerning measurements done by a colleague, he argued that not much could be concluded from too few measurements per stimulus, which deviated from his method of average error (*2*, Vol. 1, p209).

A closer inspection of Vierordt’s experiments shows various differences to the method proposed by Fechner. First of all, while Fechner was mainly interested in the just noticeable differences (JND), Vierordt reported extensively on the “constant errors”, which Fechner mentioned but treated more as a side note. Other diverging aspects of Vierordt’s method, already being criticized by Müller (*5*), was the missing temporal exchange between N and F, or the unidirectional change of the test duration (necessarily always starting from small values). At least Vierordt partly knew that his method deviated from the one Fechner had proposed, but he defended those differences by claiming several advantages (*1*, e.g., p29ff and p35). However, what is easily overlooked is that according to Vierordt in the experiments “the assistant provided … a time interval of arbitrary magnitude” (*1*, p35). According to Fechner’s and Müller’s descriptions, the method requires equal or ordered, rather than arbitrary magnitudes. Thus, evidently, the method used by Vierordt was not at all what Fechner had in mind.

Other researchers in the late 19^th^ century confirmed Vierordt’s findings (see *6*, chapter XV, for a summary), even though already Fechner criticized their results, but mostly with respect to Weber’s law (*7*). About 40 years later, Woodrow aimed to replicate Vierordt’s results but found no evidence for consistent over- and under-estimation in reproduced durations (*8*). Inspection of his methods shows that only one single interval was tested per day (50 repetitions). The author explicitly mentioned: “Entirely different results might be expected from an experiment in which the various intervals were all employed on one day, particularly if they were used in an irregular order” (*8*). Thus, presenting the stimuli one by one and with sufficient temporal separation, as suggested by Fechner, apparently avoids the systematic errors that are the characteristic of Vierordt’s law. In other words, Vierordt’s law seems to be a consequence of the particular experimental protocol.

Over the next 80 years, various other investigations followed, but without providing a formal theory for Vierordt’s law. In other fields of psychophysics and experimental psychology, effects analogous to Vierordt’s law were discovered for other types of magnitude estimation, such as “the law of central tendency” (*9*) and the “regression effect” (*10*), and were also related to sequential or serial dependence (*11*–*13*). Interestingly, Hollingworth, who also referred to Vierordt’s work, already provided important cornerstones of the effect, such as the indifference point depending on the range of stimuli given: “in all estimates of stimuli belonging to a given range or group we tend to form our judgments around the median value of the series” (*9*). He concluded these remarkable insights from a series of experiments that he published in 1909, where he compared magnitude reproduction for different ranges of stimuli and for single stimuli presented in isolation (*14*). Hollingworth’s conclusions are thus providing evidence for the importance of the context of other stimuli in which a particular test stimulus is judged.

Thus, even though the basic ideas employed later in a formal theory of the Vierordt’s law and the central tendency (for review see *15*–*17*) have been laid out early on (*8*,*9*), explanations for these and other related phenomena, such as the range effect, have been scarce. In experimental psychology and related fields, most studies recognize and accept those types of systematic errors as a usual finding. To our knowledge, the first study offering a quantitative formal theoretical treatment of the central tendency based on prior expectations was published at the end of the 20^th^ century but has been completely overlooked by the scientific community (*18*). Interest was, however, revived a couple of years ago by the Bayesian approach for perception (e.g., *19*). We independently proposed a theory of central tendency and range effects for magnitude estimation based on iterative Bayesian inference combined with the Weber-Fechner law (*16*,*20*,*21*), which offered a concise explanation for the central tendency, range and order effects, and quantitatively showed how prior information on stimulus range was updated during the course of the experiment. In parallel, Jazayeri & Shadlen (*22*) also proposed that the central tendency is an outcome based on integrating current sensory input with prior information about the range of the stimuli. Several similar modeling efforts followed (*23*–*26*), but none took a closer look at the original data.

We hypothesized that if 1) Vierordt’s law is a consequence of the experimental protocol, and 2) iterative Bayesian estimation can explain the central tendency, then we should be able to predict Vierordt’s original data using our model (see *Supplementary materials SM 1*) by applying the original experimental protocol as closely as possible. We simulated the 1104 randomized stimuli from 22 intervals described in Vierordt’s Table A. Since the exact magnitude and order of stimuli is not given, we repeated the simulation 10000 times (each time fitting the single parameter of the model to Vierordt’s data) and selected the best result (see methods in *SM 2*). The simulation did not include sensory or motor noise (as in *21*). Figure 1A depicts the Vierordt’s original data together with the best fit from the simulation, and Figure 1B shows the best simulated sequence.

Evidently, the model provides an excellent fit to the Vierordt’s data. However, how much does the reproduction error depend on experimental protocols? Supposing the same intervals are provided in ascending or descending order (assuming the same model with identical parameters), the model predicts the absolute percentage error would be below 0.2% for all intervals (as compared to below 15% in the original Vierordt’s data). The differential outputs of the simulation corroborate our suspicion that Vierordt’s law is a consequence of the random presentation of stimuli within the same experimental context. The iterative Bayesian updating model thus can explain both Hollingworth’s conclusions about the central tendency and Woodrow’s failure replication of Vierordt’s findings.

However, repeatedly presenting the same stimulus or using strictly ascending/descending series of stimuli may introduce other types of errors, such as habituation and expectation errors. Thus, the question arises whether we have to abandon randomized stimulus presentation at all or whether we have to tolerate the range-dependent systematic errors. The answer is a direct consequence of our iterative Bayesian model, which has an important underlying assumption, namely the magnitude/intensity of the stimuli out there in the world mostly remain constant except for a small random change. The model assumes that the change of the magnitude follows a Wiener process (random walk) from one trial to the next. If this assumption is satisfied, the model provides an optimal estimator for the slowly changing stimulus magnitude. Hence, if the sequence of the stimuli follows the properties of a random walk, the central tendency should be suppressed compared to that of a randomized sequence. Figure 1C illustrates this difference between a random walk sequence (Figure 1D) and a randomized sequence (note that stimuli in 1B and 1D are the same except for the temporal order of presentation).

To empirically validate this prediction, we conducted a duration reproduction experiment (see details in *SM 3*), in which 400 randomly selected stimulus durations (from 0.4 to 1.9 seconds) were tested in two sequences: 1) the stimulus order conformed to a random walk, and 2) the randomized order. We fitted a linear regression individually for each participant to the reproduced duration plotted over the test duration and used (1-*slope*) as regression index (see *SM 1* for the connection between model parameter and regression index). A regression index of 0 corresponds to no central tendency, while an index close to 1 shows a strong central tendency effect.

A repeated measures ANOVA of the regression indices revealed a significant effect of randomization [F(27,1)=53.5, p<0.0001] with the average index close to 0 (mean±SD 0.095±0.138) for the random walk, but much higher regression index (0.456±0.173) for the randomized sequence (see supplementary Figure S1 for individual data). Figure 2A shows the percent error of reproduction as a function of the test duration for both conditions. While Vierordt’s law is still visible for the “random walk” sequence, the average errors are drastically lower than those in the randomized sequence, despite both conditions containing the same set of stimuli, but presented in a different order. We also fitted the dynamic-updating Bayesian model to the reproduction data from the “randomized” condition for each participant separately and used the fitted model parameter to predict the results from the “random walk” (predicted average regression index 0.011±0.011). The predicted results are shown in Figure 2B. A comparison of actual and predicted data shows that the average central tendency in the “random walk” condition is higher than predicted from the model simulation. Closer inspection shows that the learning rate for the prior in the “random walk” condition is, at least for some participants, slower than expected from the model fitted to the “randomized” data. However, overall there is a good correspondence between data and prediction: the central tendency does not vanish but becomes significantly smaller.

**Fig. 2.**
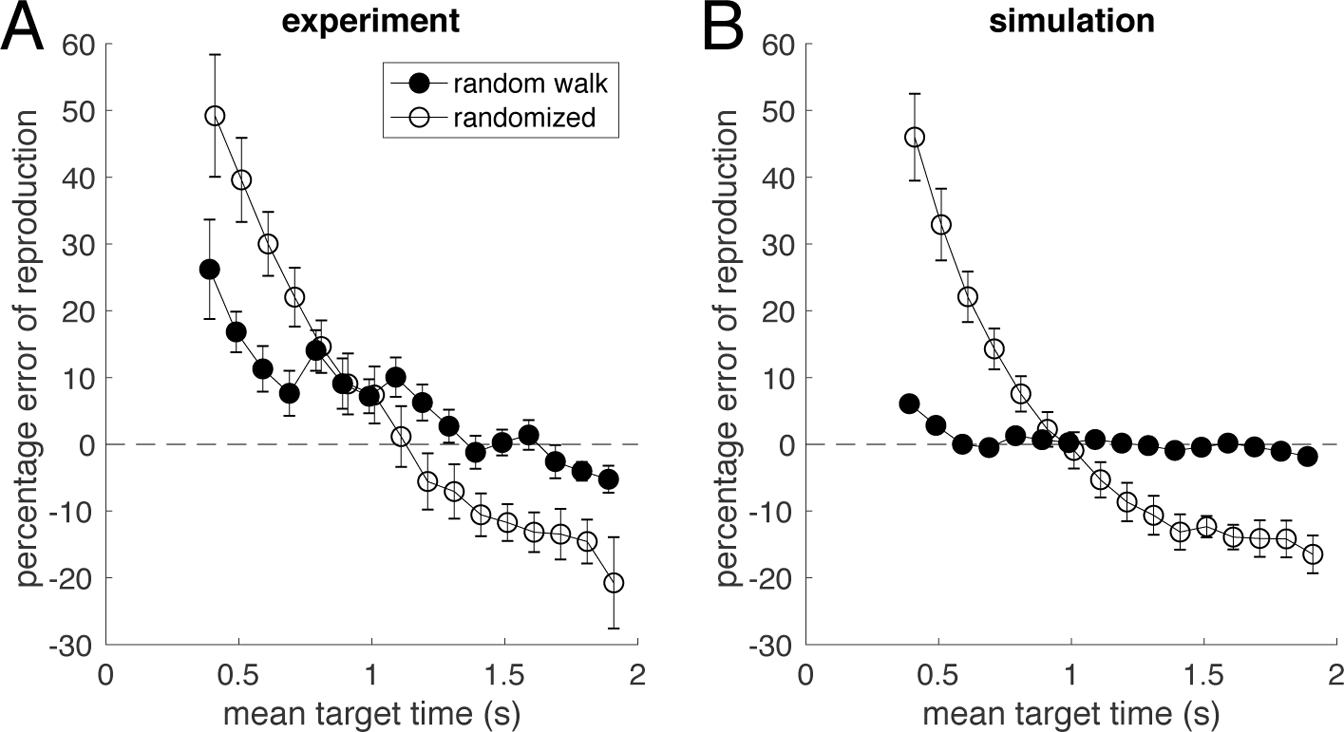
Reproduction experiment with stimuli ranging from 0.4s to 1.9s (400 durations per subject per condition). **(A)** Average reproduction error as a function of the duration from 14 subjects in two experimental conditions (open circles: randomized, closed circles: random walk). **(B)** The simulation of the iterative Bayesian model for the “randomized” data (open circles, best fit for each participant separately) and prediction for “random walk” data (closed circles, using the individual model parameters from the “randomized” experiment). Error bars show standard error of the mean.

Both data and model simulation thus confirm our hypothesis that the major factor for Vierordt’s law is the experimental protocol. While other factors still play a role - especially the range of stimuli presented - systematic errors can thus be minimized by an appropriate stimulus design even without abandoning randomization completely, but instead resorting to random walks. Notably, our experiment also extends Hollingworth’s claims (*9*,*14*): it’s not just the range of magnitudes presented that determines the central tendency, but even more so their order. Moreover, our findings refute models that assume a *static* prior distribution depending on the range of stimuli as reason for the central tendency (e.g., *22*). In such a case, we would expect similar results for both conditions, since both the range and the stimuli are identical.

In summary, from a re-evaluation of the original dataset with iterative Bayesian modeling, and validation by new experiments we conclude that Vierordt’s law (and the central tendency) is a result of the specific experimental protocol - randomly presenting stimuli with large trial-to-trial magnitude fluctuation. This protocol deviates from what usually happens in everyday life, where either successive magnitudes are equal and share the same context, or different magnitudes are associated with different contexts. The proposed underlying mechanism of Bayesian dynamic updating indeed improves performance over trials for equal or slowly changing magnitudes but not for large magnitude fluctuations. According to our analysis, 150 years of research on Vierordt’s law have thus focused on an effect that is caused by an unnatural but since then widely adopted experimental protocol, which was first introduced by Vierordt, who misinterpreted the method of reproduction invented by Fechner and described in his groundbreaking “Elemente der Psychophysik” (*2*)^1^. After all, it was not Fechner’s fault.

## Footnotes

1 Notably, Fechner stated (*2*, Vol. 1, p 84, translation by SG): … if one makes this or that number of observations, once in this sequence, then in another, under these or those circumstances without fixed rules, then the usefulness of the observations suffers in every respect.

## Acknowledgments

**Funding:** This study was supported by German DFG research projects SH 166/3-2 and GL 342/3-2. We thank Xuelian Zang and Xiuna Zhu for collecting behavioral data.

**Author contributions:** SG modeled the original Vierordt’s data, ZS conducted the validation experiment, and SG and ZS wrote the manuscript.

